# Developmental transformations of Purkinje cells tracked by DNA electrokinetic mobility

**DOI:** 10.1101/2024.08.29.610366

**Authors:** C. Brandenburg, G.W. Crutcher, A.J. Romanowski, S.G. Donofrio, L.R. Duraine, R.N.A. Owusu-Mensah, I. Sugihara, G.J. Blatt, R.V. Sillitoe, A. Poulopoulos

## Abstract

Brain development relies on orchestrated placement and timing of neurogenesis in progenitor zones to produce the expansive cellular diversity of the brain. We took advantage of bioelectric interactions between DNA and embryonic tissue to perform “stereo-tracking”, a developmental targeting strategy that differentially labels cells positioned at different depths within intact progenitor zones. This three-dimensional labeling was achieved by delivery of plasmids with distinct electrokinetic mobilities into neural progenitor zones *in utero*. We applied stereo-tracking with light sheet imaging in the cerebellum and identified that Purkinje cells follow embryonically committed developmental trajectories linking distinct progenitor zone fields to the topography of the mature cerebellar cortex. In the process of stereo-tracking, we identified a previously unreported subcellular structure on the axon initial segment of Purkinje cells. These structures, we termed “axon bubbles”, are developmentally timed and differentially labeled by lipid-modified proteins. Our findings demonstrate key rules that orchestrate the stereotyped transformations from fetal progenitors into mature networks of neuronal circuits, and demonstrate the potential of progenitor zone stereo-tracking to reveal new biology within intact developing systems.

## INTRODUCTION

Neurodevelopment unfolds with a complex orchestration of cellular transformations from mitotic progenitor zones that line the ventricles, to grey matter areas that harbor the cell bodies of postmitotic neurons embedded in mature neural circuitry. Tracking the stereotyped patterns of these transformations is an important aspect of developmental neuroscience, and is enabled by powerful molecular methods, such as *in utero* DNA electroporation^1^ and lineage tracing^2^. Despite these advances, brain areas with complex patterning, like the cerebellum, still present significant challenges in revealing their developmental programs.

Purkinje cells, central to cerebellar circuits, have a complicated molecular program and migration trajectory that is difficult to track across development because of the transient nature of gene expression and the formation of clusters before dispersal into the expanding foliated lobules^3–5^. Despite this complexity, their ultimate placement within the mature cerebellum appears to be determined at neuron birth^6–8^.

Here, we identify a bioelectric interaction between DNA and neural progenitor zones, and leverage it to track neuronal trajectories as they migrate from distinct progenitor zone fields to their positions in the cerebral and cerebellar cortices. We show, through morphological analyses of intact cerebella in 3D, that detailed topographic information across the Purkinje cell layer is determined embryonically through local positioning in progenitor fields. In this process, we additionally uncovered previously undocumented subcellular structures emerging during Purkinje cell development that we term “axon bubbles”. Finally, we provide technical details and resources for investigations into the development of neuronal topography using a labeling toolkit for progenitor zone stereo-tracking and 3D analysis.

## RESULTS

### Differential plasmid DNA expression based on bioelectric interactions *in utero*

We observed that mixes of plasmids electroporated *in utero* can be expressed in distinct daughter-cell populations of a neural progenitor zone, even though expression is driven by the same promotor. We investigated the circumstances under which plasmids from a single electroporation pulse (Fig. 1a) appear to segregate into distinct neurons by testing combinations of plasmids ubiquitously expressing green and red fluorescent protein variants (GFP and RFP; Extended Data Fig. 1). We determined that segregation of mixed plasmids into distinct postmitotic neuron populations was not driven by promoter or plasmid sequence, but rather was an electrogenic effect correlating with the relative electrokinetic mobility (EKM) of the plasmids in the mix. Similar to how EKM determines plasmid electrophoretic mobility and separation of DNA mixes on a gel according to molecular weight and supercoiled state^9,10^ (Extended Data Fig. 2), we observed plasmid expression segregating into distinct daughter-cell populations according to each plasmid’s EKM (Fig. 1b-e).

**Figure 1:**
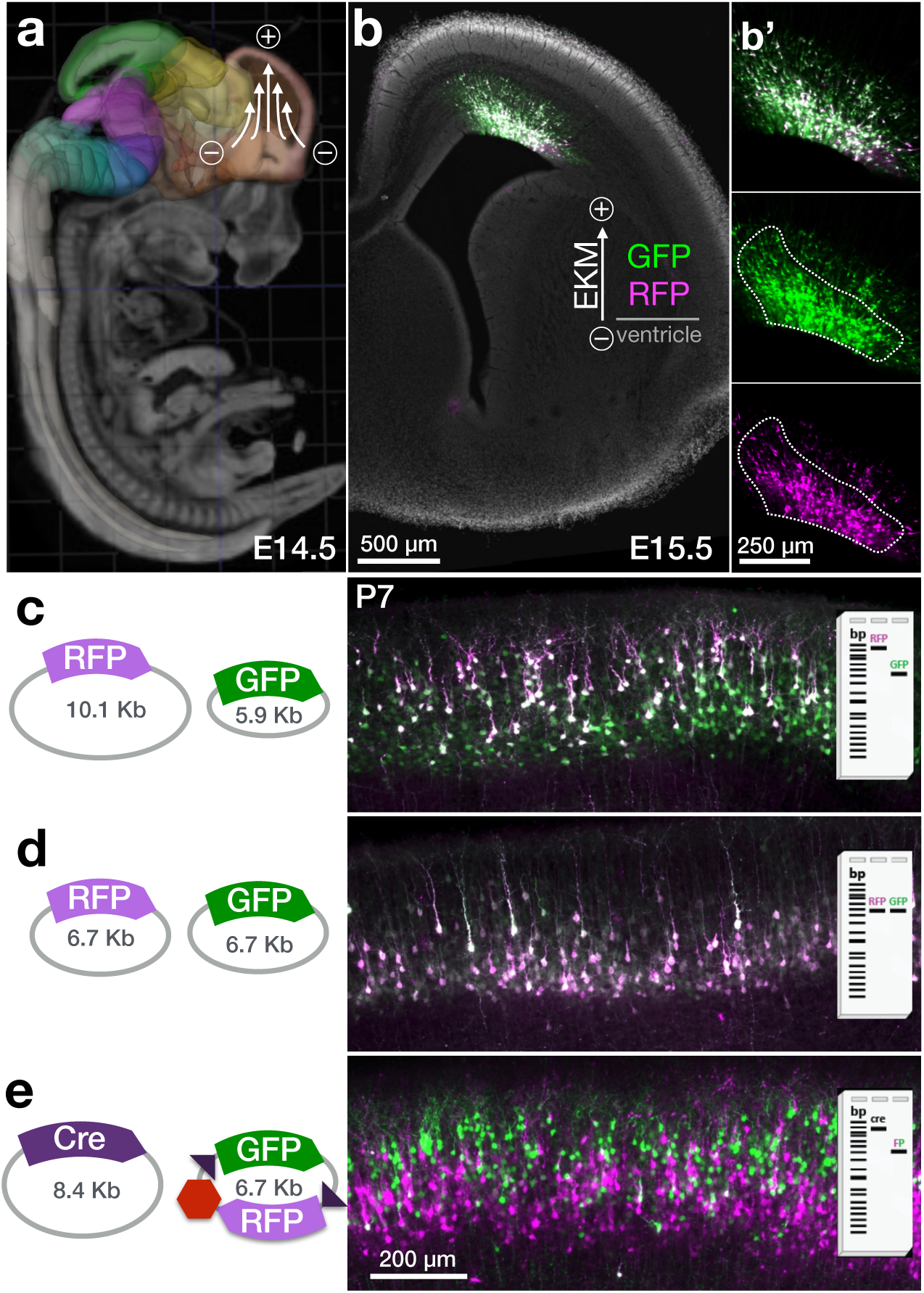
Segregation of plasmids by electrokinetic mobility within neural progenitor zones. **a,** Schematic of tri-pole electrode (tritrode) *in utero* electroporation in the developing cerebral cortex (orange, from Allen Developing Brain Atlas). DNA injected in the lateral ventricle is targeted by a focused electric field into neural progenitor cells of the pallial neurogenic zone at the dorsal surface of the ventricle. Plasmid electroporation at E14.5 leads to fluorescent protein expression in migrating postmitotic neurons as seen 24h later (**b,b’**). A temporal range of neurons can be seen among those electroporated, representing a front end of cells with a head-start in migration. At E15.5, neurons with the highest expression are closest to the ventricular surface (white outline in **b’** inset), at the shortest distances DNA is required to travel. Those cells receive the highest copy number of both plasmids. Expression rapidly declines with distance from the ventricular surface, and more so for larger plasmids (magenta) with lower electrokinetic mobility. As such, cells with a head start in migration that are farther from the ventricular surface preferentially receive plasmids with higher electrokinetic mobility. **c,** Simultaneous electroporation of a mixture of plasmids of uneven size (schematized as gel bands with different electrokinetic mobility) at E14.5 results in plasmids expressed in neurons at distinct radial depths at P7. Larger plasmids are expressed in more superficial neurons (late-born, magenta), while smaller plasmids are in both superficial and deeper layers (early-born, green). **d,** Size-matched GFP and RFP plasmids (6.7 Kb) express homogeneously in the same neurons. **e,** Mixture of a large plasmid encoding Cre (8.4 Kb) and a small (6.7 Kb) bicistronic plasmid encoding Cre-ON GFP (green) and Cre-OFF RFP (magenta), shows that the majority of cells get the small plasmid (magenta and green cells), while only more superficial neurons receive the larger Cre plasmid (green cells). Cells with a head-start in migration are farther from the ventricular surface at the time of electroporation, are thus less likely to receive larger and electrokinetically less mobile plasmids, and occupy deeper layers of cerebral cortex as determined by inside-out radial migration. Thus, electrokinetically distinct plasmids can differentially label early– vs late-born neurons from a progenitor zone. Full plasmid schemes are provided in Extended Data Fig. 1. Scale bars as indicated.

When two plasmids of equal EKM (the same size and supercoiled state) were co-electroporated into the progenitor zone of the dorsal pallium, they co-expressed in the same postmitotic daughter neurons in the cortical plate (Fig. 1d). However, when electrokinetically faster and slower plasmids (lower and higher molecular weights, respectively) were co-electroporated, the faster plasmid was additionally expressed in a separate fraction of postmitotic neurons, which did not receive the slower plasmid. Neurons that received only the faster plasmid were consistently farther along in radial migration at embryonic day 15.5 (E15.5) (Fig. 1b, b’) and resided in deeper layers at postnatal day 7 (P7) than neurons that received both faster and slower plasmids (Fig. 1c). Given the inside-out progression of development in the cerebral cortex^11,12^, these indicate that faster plasmids reached an earlier-born population of neurons, which had a head-start in radial migration, and ended up in deeper layers of the cortical plate.

We interpret these observations as follows: the electroporation pulse at E14.5 caused a bioelectric interaction in which each plasmid traveled a distance within the progenitor zone proportional to its EKM. Within the given duration of the electric pulse, faster (smaller and/or supercoiled) plasmids penetrate deeper into the progenitor zone compared to slower (larger and/or relaxed) plasmids, much like plasmid migration in gel electrophoresis. Thus, the faster plasmid of the mix reaches deeper fields of the progenitor zone, targeting older cells that are farther along in the developmental progression of neurogenesis, interkinetic nuclear migration, and radial migration.

We cannot discern the stage in the developmental continuum of neurogenesis, interkinetic nuclear migration, and radial migration in which plasmids are received, however there appears to be a range that is bisected by our approach between the slowest plasmids targeting progenitors on the ventricular surface that correspond to labeled postmitotic neurons in the most superficial layers of the cortex, and the fastest plasmids targeting older cells on their way into radial migration corresponding to labeled postmitotic neurons in deeper layers. This range appears to be analog, since the separation of plasmid expression in the layers of cortex, as well as the intensity of fluorescence scales depending on the EKM difference between the two plasmids and the titrated concentrations (Extended Data Fig. 3).

### Manipulation of DNA electrokinetic mobility enables stereo-tracking in neural progenitor zones

We leveraged the differential targeting of daughter-cell populations with a single electroporation pulse with plasmids of distinct EKMs to establish an *in utero* labeling approach for differential labeling of cells that are superficial vs. deeper into ventricular progenitor zones of the brain. We term this approach “stereo-tracking” as it enables simultaneous labeling of cells at distinct depths within a 3-dimensional progenitor zone.

In the radially organized progenitor zone of the dorsal pallium, shallow– vs. deep-labeled cells correspond to younger vs. older, respectively, postmitotic neurons within the progenitor zone at the time of the electroporation pulse. During the ensuing “chase” period, the trajectories of two generations of postmitotic neurons from the same progenitor zone can be tracked to their distinct layer positions in the cortical plate (Fig. 1, Extended Data Fig. 3).

Stereo-tracking can be viewed as a developmentally-orthogonal labeling strategy to lineage tracing^2^. While lineage tracing vertically labels all generations of cells from an individual progenitor, stereo-tracking labels a snap-shot of distinct cell populations based on their local fields within the volume of the progenitor zone. This includes labeling younger vs. older generations, most prominent in simple progenitor zones, as well as convolutions of internal 3D topography in more complex progenitor zones, as in the example of the cerebellum below.

To facilitate labeling clarity of younger vs. older batches of postmitotic neurons, we employed a binary labeling strategy based on Cre recombinase, similar to those previously employed for *in utero* mosaic manipulation approaches^13,14^. The faster plasmid encodes floxed-RFP-stop GFP and the slower plasmid encodes Cre, such that older daughter cells receive the faster plasmid and express RFP alone, while younger daughter cells receive both faster and slower plasmids, suppressing RFP and expressing GFP alone (Fig. 1d). With this conditional regime, we constructed plasmids with high expression of bright variants of GFP and RFP, as well as optional modifications to facilitate membrane labeling (See methods and Extended Data Fig. 1).

Using bilateral tritrode electroporations, we titrated the amounts of plasmids to approach labeling parity in younger and older populations and to achieve the highest FP expression, both of which were affected by Cre concentrations (Extended Data Fig. 3). A further internal gradation dependent on postmitotic cell age can be observed within each color, correlating broadly with fluorescence intensity. While intensity also varies stochastically from one cell to the other based on plasmid copy number received, at the population level, intensity scales with age within each color population, such that brighter cells are generally younger than dimmer cells of the same color (Extended Data Fig. 3).

With these advances, progenitor zone stereo-tracking can achieve reproducible, staggered labeling of superficial vs. deeper cells in a ventricular neurogenic zone with a single electroporation pulse, revealing during chase periods their migratory paths and topographic relationships as they integrate into mature circuitry. This strategy offers the opportunity to investigate how the spatial organization of progenitor zones contributes to patterning in brain areas with poorly-understood development.

### Progenitor zone stereo-tracking in the cerebellum reveals topographic migration of Purkinje cells

We employed stereo-tracking to investigate the development of Purkinje cell layer migration and topography in the cerebellum. Purkinje cells are born within a narrow three-day window (E10.5-E12.5) in a compact progenitor zone above the 4^th^ ventricle^15–18^ (Extended Data Fig. 4). After neurogenesis, they migrate into the Purkinje plate^11,19–21^, undergo a period of clustering, and then disperse from the centralized cluster stage postnatally (P3-P10)^7,22^ into the foliating lobules and into their characteristic pleated monolayer within the mature cerebellum.

We asked whether the specific timing of neurogenesis of Purkinje cells contributes to their topographic patterning in the cerebellar cortex. To do so, we used stereo-tracking to segment nascent Purkinje cells with a pulse at E11.5 (Fig. 2a) bisecting the neurogenic window into early– and late-born Purkinje cells across the anteroposterior axis. Using the conditional strategy of Fig. 1d, EKM-discrepant plasmid mix was injected into the 4th ventricle and electroporated at E11.5 to label deep progenitor zone cells with RFP (magenta) and shallow cells with GFP (green, Fig. 2b-i). Stereoscope imaging allowed for examination of the overall distribution of Purkinje cells at P14, by which time they have migrated into their final positions within the cerebellar cortex.

**Figure 2.**
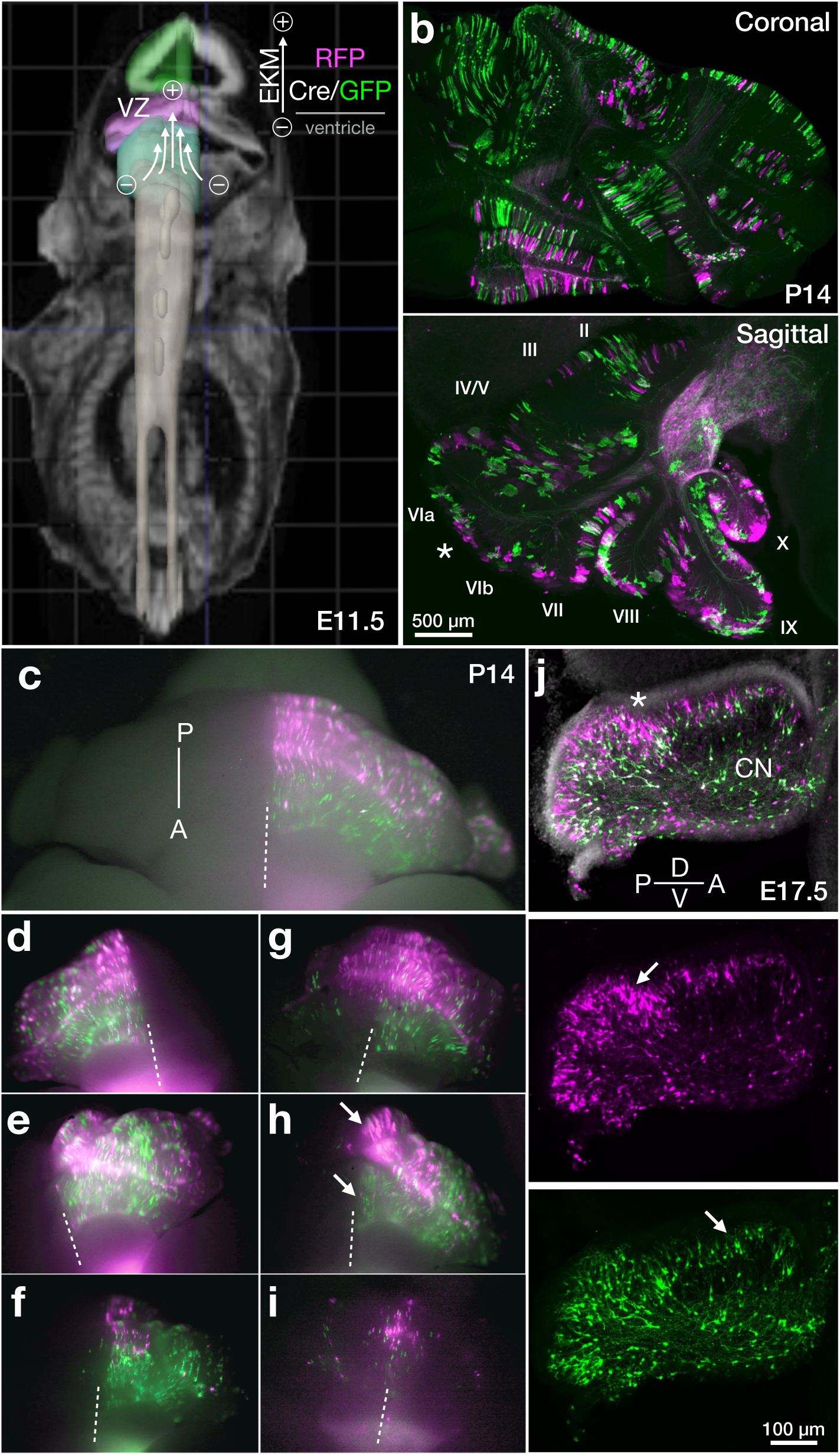
Stereo-tracking Purkinje cells in the developing cerebellum. **a,** Tri-pole electrode (tritrode) electroporation schematic to target Purkinje cells in the ventricular zone superior to the 4^th^ ventricle at E11.5 (Allen Developing Brain Atlas). Stereo-tracking plasmid pairs used throughout the figure (Extended Data Fig. 1b) label deep cells with GPI-RFP (magenta) and shallow cells with Cre/GPI-GFP (green) as indicated in the EKM schematic. **b,** Coronal and sagittal sections at P14 produce complex patterns of stereo-tracked Purkinje cells that are difficult to appreciate in 2D sections. **c,** Epifluorescence stereoscope overview of whole uncleared cerebella, Purkinje cell electroporations, mostly on one cerebellar hemisphere (except g and i). Clear segregation of stereo-tracked cells can be seen across electroporations, particularly at the sagittal level of the vermis. Anterior-posterior orientation is indicated (A-P). The midline is indicated with a dashed line. With a strong electrical field, highest plasmid expression includes most lateral portions of the flocculus. **d-i,** The strength of the electrical field and exact embryo age in hours may affect the outcome of electroporation. However, the posterior vermis consistently shows a high proportion of deep-labeled RFP cells (magenta), revealing that individual Purkinje cells have migration patterns that are committed and represented in their progenitor zone position already by E11.5. Shallow-labeled Cre/GFP (green) cells are predominant in anterior lobules, while the deep-labeled RFP cells (magenta) are predominant in posterior lobules. **j,** At E17.5, there is distinct layering of the two stereo-tracked Purkinje cells of the posterior lobule, similar to that seen in the cerebral cortex. Anterior lobules have less Purkinje cell layering. The asterisks in b and j correspond to the separation of posterior from anterior lobules, arrows indicate the bright RFP cluster at E17.5 and its corresponding location at P14 and the anterior layers that are majority GFP positive at E17.5 compared to P14. Scale bars as indicated.

We observed a striking dispersal of generation-tracked Purkinje cells across entire lobules at P14, with consistent topographic segregation. RFP and GFP are both represented in the posterior cerebellum (Fig. 2b sagittal), with the highest density of RFP-expressing cells in the vermis, while GFP dominated the anterior lobules (Fig. 2c-i). This pattern was reproducible across different electroporations.

The challenges of targeting the Purkinje cell progenitor zone, due to the early embryonic stage and central location^9,23^, result in varying levels of electroporation efficiency across individual electroporations, with the larger Cre plasmid more variable due to its larger size. Through this intrinsic variability, we observed that in weaker electroporations, and/or when fertilization took place toward the later end of the overnight breeding window, the parasagittal stripe pattern previously reported to be dependent on Purkinje cell birthdate^4,5^ becomes evident by stereo-tracking (Fig. 2i). For example, the weak electroporation in Fig. 2i leads to expression in only late-born cells at the ventricular surface, which corresponds to the parasagittal stripe pattern observed after E12.5 injection of birth-dated adenoviral vector expression^5^, which then bisects the late-born cells into the two color labeling at that time point. Stronger electroporations delivered precisely at E11.5 label across parasagittal clusters and stripes as cerebellar compartmentalization matures (Fig. 2c-h).

In addition to that pattern, when examining the high-intensity cells within one color, which as mentioned above, as a population correspond to overall deeper cells at the time of the pulse, we observed that one area of the vermis always has the brightest RFP cells (Fig. 2h upper arrow, Fig. 2j, arrow at superficial cells separating the posterior from anterior progenitor zone), indicating that not only are stripe locations based on birthdate, but that topography within the stripes also correlate with Purkinje cell progenitor location.

When the electrical field is strong (e.g. Fig. 2h), the posterior vermis contains the highest RFP expression, while the anterior regions are dimly GFP positive. This is due to the anterior region of the progenitor zone being intrinsically shallower (or having fewer layers, defined as the numbers of Purkinje cells that are stacked in the dorsal-to-ventral axis) than the posterior region (Fig. 2j, Extended Data Fig. 4,24^−26^), which biases toward GFP expression being closer to the ventricular surface.

As in the cerebral cortex (Fig. 1b), it appears that cells several layers deep are electroporated within the progenitor zone, with those closer to the ventricular surface receiving the highest copy number of FP plasmids and therefore become the brightest. However, when a larger Cre plasmid is introduced, the layers closest to the ventricular surface receive more copies of Cre, turning the most superficial cells at the ventricular surface into Cre-ON GFP expressing, superficial labeled, cells.

At E17.5 (Fig. 2j), this differential layering based on plasmid expression can be seen, similar to that observed in the cortex (Fig. 1d). However, this apparent layering disappears by P7, as the Purkinje cells originating closest to the ventricular surface occupy the same monolayer as the bright RFP expressing cells that originated from a deeper field of the progenitor zone. Therefore, from the time of the electroporation pulse (∼E11.5) to ∼P7 when the Purkinje cells form a monolayer, the overarching pattern in the cerebellum resembles the organization of the cortex, in that superficial cells closely follow deeper cells^7,8^ as they migrate away from the posterior ventricular zone.

The anterior cerebellum must also accommodate a substantial number of cerebellar nuclei neurons. Therefore, layering within clusters is thinner than the more caudal aspects of the posterior zone^26^ (Extended data Fig. 4), which translates to most cells becoming GFP positive. The asterisks in Fig. 2b and 2j, highlight the same strongly electroporated posterior lobule cells at E17.5 and P14. The arrows in Fig. 2h and 2j additionally highlight how the brightest cells near the border of posterior and anterior lobules are consistently electroporated, which translates to one location in the posterior vermis at P14 across electroporations (Fig. 2c-i) when examining with stereoscope images. Therefore, individual Purkinje cells do not disperse randomly from the cluster stage into the monolayer. Rather, Purkinje cells of the same progenitor zone field are embryonically committed to follow a stereotyped trajectory that consistently leads them to the same topographic location within a lobule in the adult cerebellar cortex.

The overall stereo-tracked electroporation patterns are summarized in Fig. 3a. The shape of the Purkinje cell plate resembles a cornucopia, with more Purkinje layers in the posterior zone than the anterior zone (Extended Data Fig. 4), which can only be electroporated by the small RFP plasmid at the peak and leads to the high density of RFP cells in the vermis across electroporations. Fluctuations lowering Cre concentration or electroporation efficiency, shift gradients to more RFP cells overall (Fig. 3j); while with higher Cre fluctuations only the peak of the progenitor zone remains RFP, unreached by the large Cre plasmid (Fig. 3f).

**Figure 3.**
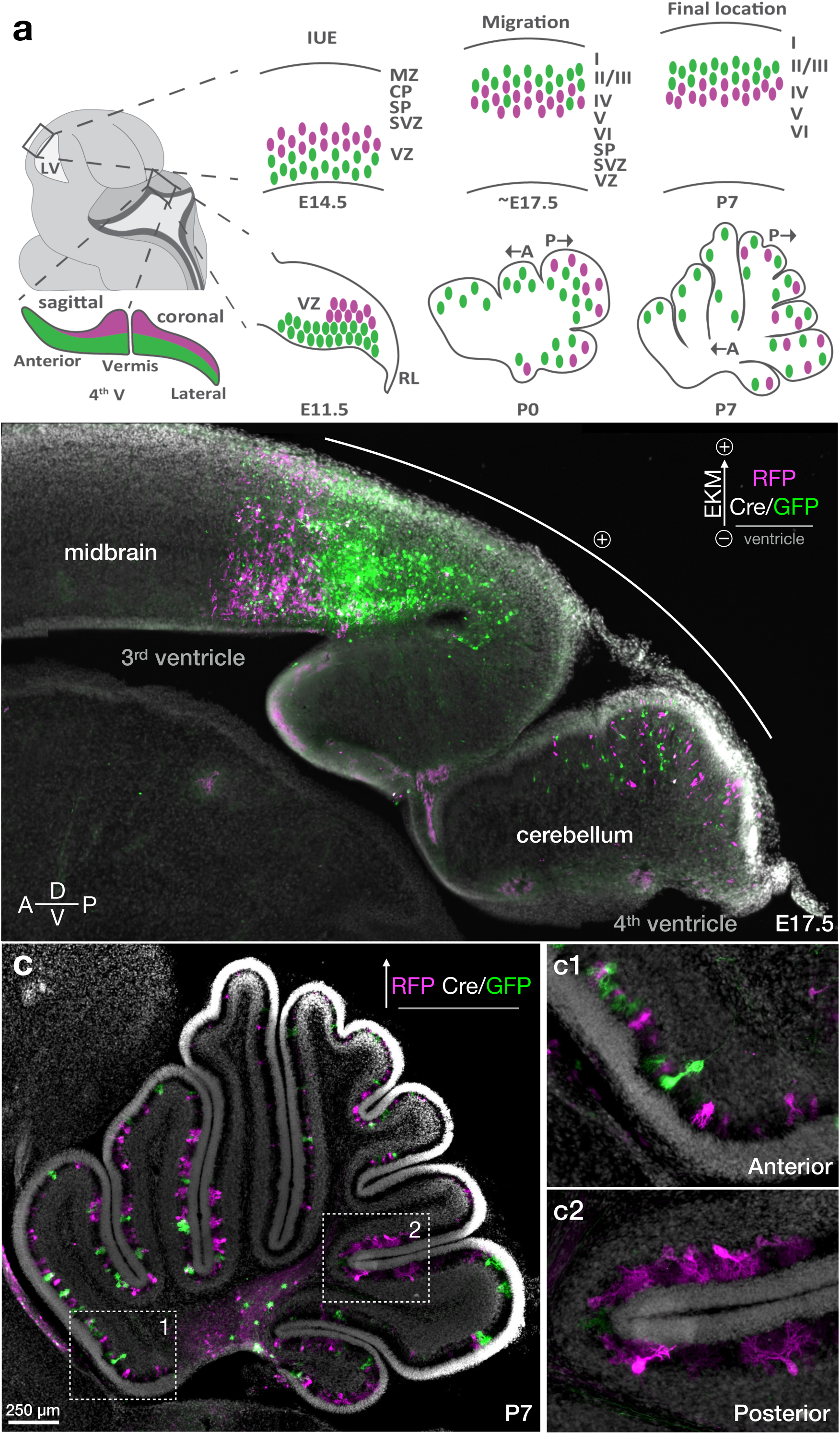
Embryonic commitment of postnatal Purkinje cell topography in the cerebellar cortex. **a,** Model of Purkinje cell development based on stereo-tracking data. The cerebellar ventricular zone (VZ) is not uniformly shaped as is the progenitor zone of the dorsal pallium (see Fig. 1), a gradient of stereo-tracked cells is seen along the anteroposterior and mediolateral axis. The anterior zone has fewer layers of Purkinje cells (see Fig. 2j, Extended Data Fig. 4), thus most cells are shallow-labeled (green) near the ventricular surface. The posterior zone has a deeper progenitor field with a higher proportion of deep-labeled cells (magenta). Purkinje cells do not migrate past earlier born cells as neurons in the cerebral cortex do. Rather they radiate outward in sequence with their position in the progenitor zone toward the outer cerebellar surfaces. Once the Purkinje cell monolayer is formed, the majority of cells in the anterior zone will derive from shallow progenitor fields (green), while the posterior zone comprises a mosaic of both deep and shallow-labeled Purkinje cells. **b,** At E11.5, embryos are small and difficult to target so there is variation in the resulting electroporation. As an example at E17.5, the bias of the electrical field toward the third ventricle resulted in plasmids strongly electroporating the midbrain and showing a distinct segregation of mismatched EKM plasmids. The anterior cerebellum buried more deeply in the tissue, was not electroporated efficiently. Ensuring placement of the positive electrode directly over the Purkinje cell progenitor zone is critical for consistent electroporations. **c,** When plasmids are EKM-matched and Cre is titrated appropriately, the anterior cerebellum has equal representation of RFP (magenta) and GFP (green). This allows for separating individual cells stochastically without biases of progenitor zone fields. A developmental gradient can be seen at P7 where anterior lobules have Purkinje cells with more immature dendritic morphologies (dashed box 1, inset **c1**), compared to those of the posterior cerebellum that have more well-developed morphology (dashed box 2, inset **c2**). Some Purkinje cells can still be found migrating along axon tracts, highlighting the dynamic nature of cerebellar development in early postnatal stages. Scale bars as indicated.

Since layering in the Purkinje cell plate is reduced in the anterior and lateral ventricular zone, the binary fluorophore effects become less pronounced outside of the vermis. Additionally, the anterior zone Purkinje cells are biased toward superficial labeling^27^, which influences their eventual zonal patterning^7,8^.

The weaker electroporation of the anterior lobules is likely due to its longer distance from the surface of the brain and therefore from the positive electrode (Fig. 3b). Depending on the characteristics of the progenitor zone within the electric field, stereo-tracking will reveal different patterns, as seen in the midbrain (Fig. 3b). With size-matched plasmids and accurately titrated Cre (Fig. 3c, Extended Data Fig. 3, 5), cells are equally labeled with RFP and GFP across the entirety of the cerebellum (Extended Data Fig. 6). Cells in the anterior lobules generally develop later morphologically than posterior lobules^21,28^, as evidenced by the size of the dendritic tree in insets of Fig. 3c, adding to the utility of this approach for comparing morphology and developmental profiles between distinct functional areas.

Overall, EKM-mismatched mosaic plasmids can label cells a few layers deep at the ventricular surface with the slower plasmid and a few layers beyond that with the faster plasmid in both cerebral and cerebellar ventricular zones. However, while the progenitor zone depth in the cortex is uniform, the cerebellum has a progenitor zone with a more complex 3D landscape, which commits migrating Purkinje cells already from E11.5 to the complex topographic patterns that they will assume in the adult cerebellar cortex.

### Optimizations for 3D imaging revealed Purkinje cell “axon bubbles” in cerebellar development

Despite the Purkinje cell layer being a monolayer, its convolutions in space due to the foliations of the cerebellar cortex make analyzing its topographic features particularly challenging through standard histological sectioning (i.e. Fig. 2b). To decipher the topographic patterns of Purkinje cells as they migrate to their final positions in the cerebellar cortex, we used tissue clearing and light sheet microscopy^29^ to document the positional relationships between stereo-tagged progenitors within the intact developing cerebellum in 3D.

We aimed to image a developmental time course of cleared stereo-tagged cerebella. To achieve sufficient tissue processing throughput and favorable optical qualities, we optimized the labeling and clearing process^30^ to maximally preserve endogenous fluorescence while enabling imaging deep into the cerebellar fissures (see methods). This enabled us to avoid the need for antibody staining, thereby dramatically shortening processing times and lowering fluorescence background noise. To better resolve individual Purkinje cell morphology, including dendritic arbors and axon projections, we produced membrane-tethered versions of stereo-tagging plasmids that express GPI-anchored FPs for outer-leaflet membrane labeling, and myristoylated FPs for inner-leaflet membrane labeling (Fig. 4a, b). With these modifications and by using the high quantum-yield variants of RFP and GFP (mScarlet-I^31^ and mGreen Lantern^32^, respectively), we were able to rapidly clear brains that maintained robust endogenous fluorescence in cell bodies and processes that withstood the delipidation process of tissue clearing and produced 3D maps of individual Purkinje cells with their processes in 3D (Movie S1).

**Figure 4.**
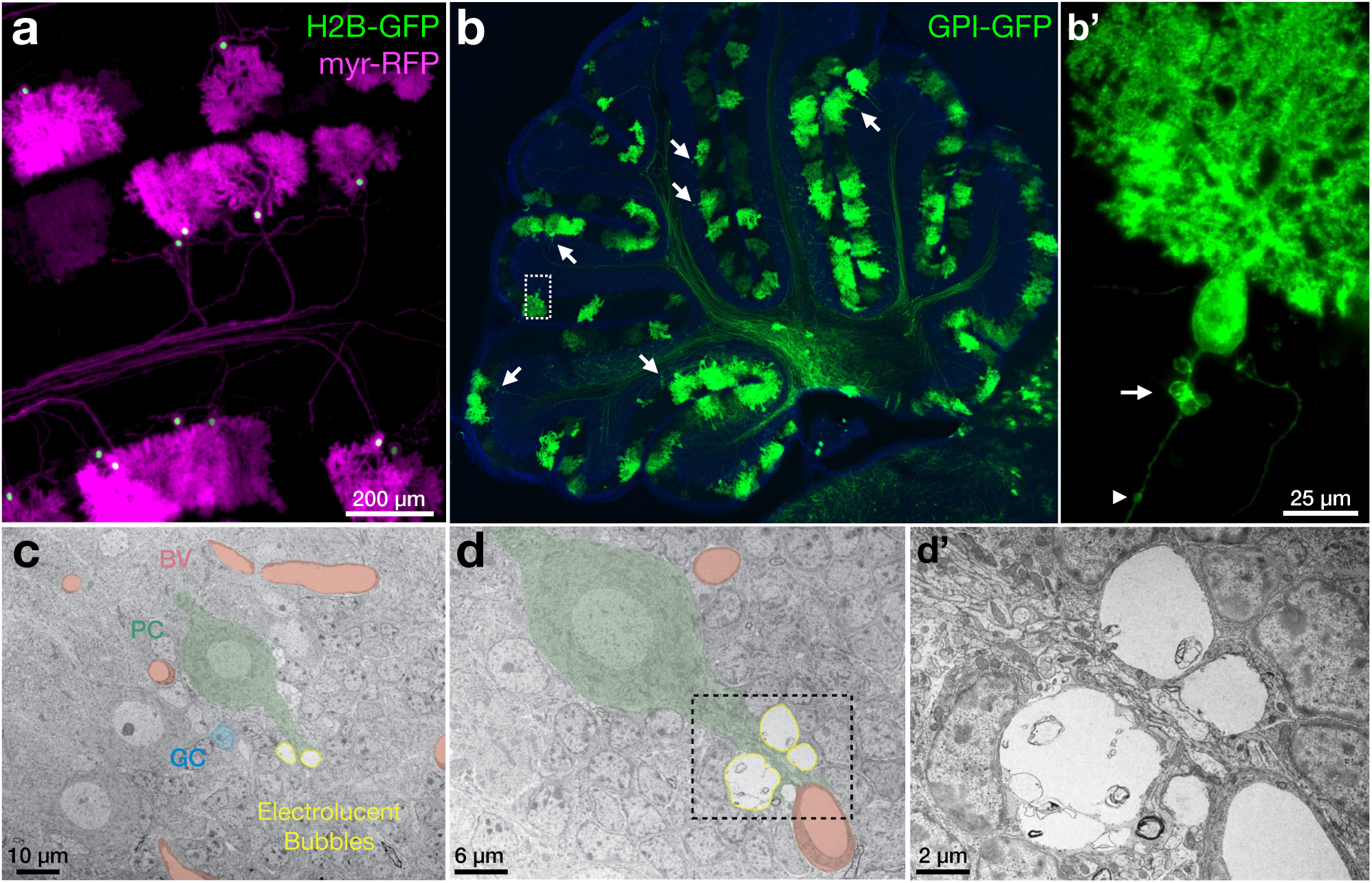
Purkinje axon bubbles are illuminated with GPI-anchored fluorophores. **a,** Electroporated Purkinje cells at P14 expressing membrane-localized myristolyated-RFP (myr-RFP, magenta) and nuclear-localized Histone 2B-GFP (H2B-GFP, green). Inner leaflet myr-RFP outlines the dendritic arbor and axons of Purkinje cells, but did not outline axon bubbles. **b,** GPI-anchored GFP (GPI-GFP, green; DAPI in blue) allows for visualization of fine processes without high fluorescence from cell bodies. Outer leaflet membrane labeling with GPI-GFP revealed Purkinje cell show axon bubbles at a consistent location at the axon hillock, approximately 10 μm from the soma at P14. Dashed square is magnified in inset **b’**, an axon torpedo-like structure is indicated (arrowhead), as well as a previously-unreported bubble structure near the axon hillock (arrow). **c-d’,** Transmission electron micrographs of unmanipulated P14 outbred strain CD1 mouse cerebellum showing electrolucent axon bubbles (yellow outline) in the precise location seen in electroporated cells at the Purkinje cell axon hillock (example shaded in green), that are unlike the blood vessels (shaded crimson) throughout the tissue, as they are not surrounded by vascular-associated cells. Numerous Granule cell nuclei (example shaded in blue) surround Purkinje cell axons at P14. **d,’** Higher magnification image of area indicated by a dashed box in d of a Purkinje cell with multiple electrolucent bubbles. Membranes can be seen to tightly associated around the bubbles with the axon membrane. Scale bars as indicated.

The distribution of lipid-modified FPs along the perimeter of neurons allowed for imaging fine processes like dendrites and axons, without being obscured by scattered fluorescence from bright soma labeling typical of cytosolic FPs, where fluorescence intensity scales with compartment volume. Unexpectedly, use of a GPI-anchored FP, specifically, revealed a previously unreported subcellular structure emerging in the postnatal stages of cerebellar development. These structures, which we term “axon bubbles”, appear by P14 at approximately ∼10 μm from the Purkinje cell soma in the likeness of a corona, near the axon hillock and pinceau (Fig. 4b’).

Axon bubbles appeared consistently in the same position at the interface of the Purkinje cell layer and the nascent granule cell layer across all areas of the cerebellum (Fig. 4b). They first appear by the second postnatal week, as they were not present at P7 in our time course, and appeared by P14. This is the phase of cerebellar development that granule cell bodies migrate *en masse* through the Purkinje cell bodies as they reposition from the superficial molecular layer to the nascent granule cell layer^33^. It is also the time when transient synapses of mossy fiber terminals onto Purkinje cell bodies and axon initial segments are eliminated to yield the mature circuitry of mossy fiber-to-granule cell innervation^34^. At P14, we see axon bubbles in roughly ∼20% of labeled Purkinje cells (Fig. 4b), suggesting that these cellular protrusions appear transiently and may be involved in the dynamic cellular processes occurring at this developmental time-point.

Cytosolic labeling has previously revealed swellings and torpedo structures in Purkinje cell axons^35–37^. Surprisingly, we did not observe labeling of axon bubbles with cytosolic FPs (Extended Data Fig 7), possibly explaining why these structures were not previously documented. Even more surprisingly, axon bubbles were also not labeled by myristoylated FPs, which are tethered on the cytosolic leaflet of cellular membranes (Fig. 4a). These observations suggest that bubbles are either an artifact caused by GPI-FP overexpression, or that the content within axon bubbles tightly excludes cytosolic components, possibly by an organelle or condensate, such that only GPI-anchored FPs on the outer or lumenal surface can be accommodated.

We turned to transmission electron microscopy (TEM) in the developing cerebellum to determine whether axon bubbles are present in native unlabeled Purkinje cells. Micrographs of outbred, unmanipulated CD1 strain mouse cerebella at P14 (Fig. 5 and Extended Data Fig. 8) confirmed the presence of structures immediately adjacent to Purkinje cell axon initial segments consistent with axon bubbles seen with electroporation of GPI-FP. These structures were observed in the corresponding location predicted by the electroporation experiments and not elsewhere within cerebellar tissue. The interior of bubbles is particularly electrolucent, consistent with exclusion of cytosolic components predicted by the lack of cytosolic FP labeling. Axon bubbles contain some inclusions that tend to be surrounded by double membranes, but do not contain recognizable axonal structures such as microtubules, neurofilaments or other matrices. This is in contrast to TEM images of axonal swellings^37^ that have clear cytoplasmic structures and are surrounded by myelin.

**Figure 5.**
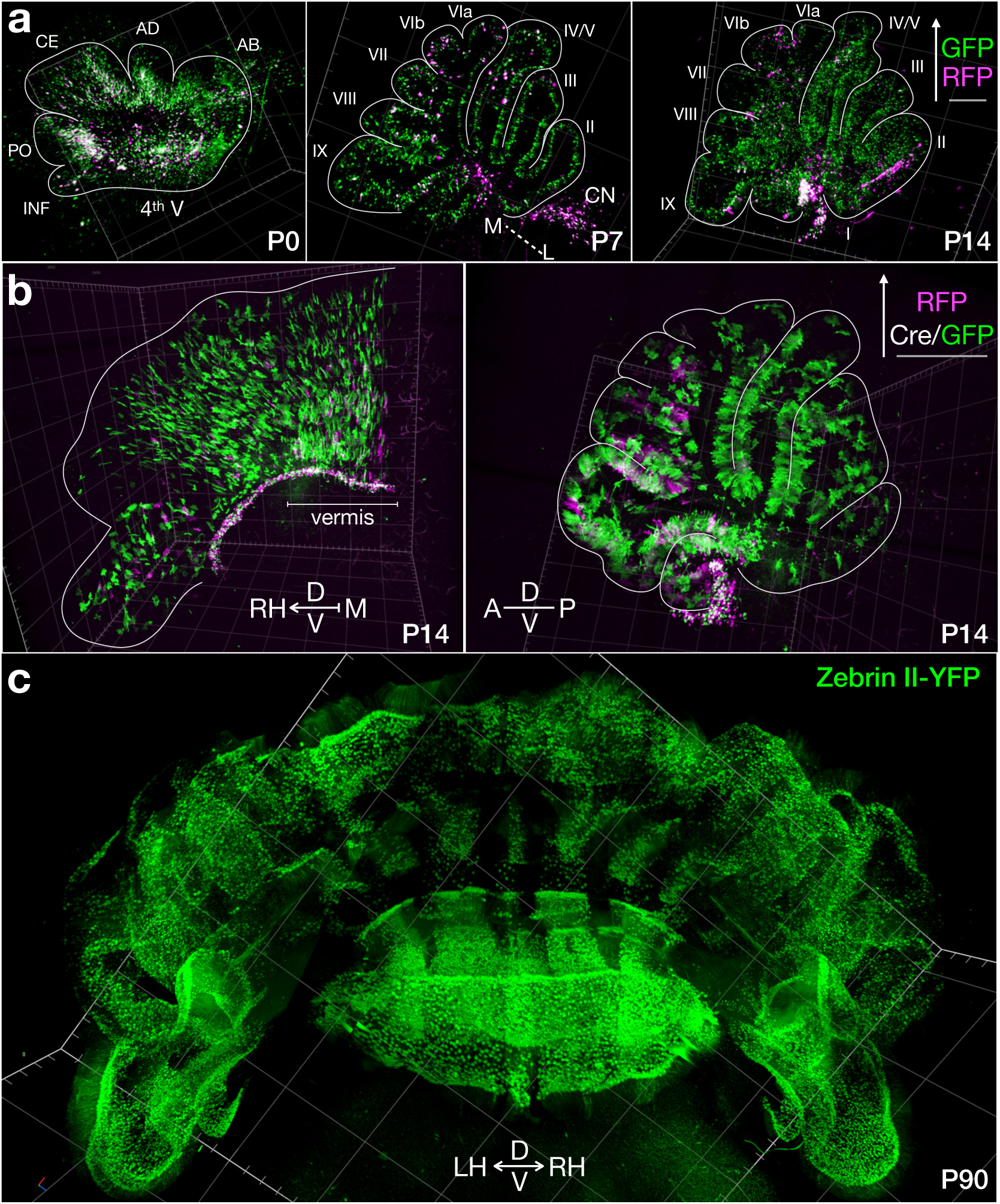
Light sheet imaging of whole cerebella after stereo-tracking across development. **a**, 3D renderings of light sheet imaging of whole cerebella after stereo-tracking with cytosolic GFP (deep, green) and RFP (shallow, magenta) plasmid pairs (Extended Data Fig. 1a) at three developmental time points as indicated. shallow-labeled RFP cells are restricted to areas of strongest electroporation. The posterior cerebellum consistently displays higher electroporation efficiency and higher proportion of deep-labeled GFP cells. One P0, one P7, and two P14 brains were rendered in 3D to identify individual Purkinje cells of each color (see Movie S2 for 3D overview). The number of cells expressing RFP was 5.8% of the number of cells expressing GFP. **b,** Whole cerebellum 3D rendering of light sheet imaging after stereo-tracking with GPI-RFP (deep) and GPI-GFP (shallow) plasmid pairs (Extended Data Fig. 1b), enabling imaging of fine processes such as dendrites and axons (see Movie S1 for 3D overview). For sagittal view, the entirety of the vermis is displayed. Purkinje cells collapse on top of each other from each plane to reveal individual lobules. 3D videos (Movies S1-2) better capture the overall distribution of labeled cells. **c,** 3D renderings of light sheet imaging of adult cerebellum from a transgenic line expressing YFP fused to aldolase C (zebrin II), processed and image with optimized clearing method, reveals contiguous and convoluted stripe topography (see Movie S3 for 3D overview).

Electrolucent bubbles are tightly surrounded by a single lipid bilayer around most of their circumference, which appears to be an extension of the Purkinje cell axolemma. There appears to be a lack of an internal lipid bilayer to separate it from other components of the axoplasm (Fig. 5 and Extended Data Fig. 8), suggesting these are not organelles of the secretory pathway, such as vacuoles. The electrolucent structures are surrounded by the Purkinje cell plasma membrane, consistent with electroporation showing GPI-FP expression delineating each bubble with narrow attachments along the axon (Fig. 4b’). We therefore surmise that these electrolucent structures represent the axon bubbles labeled by GPI-FPs in the developing cerebellum, confirming their presence in unmanipulated Purkinje cells.

With modifications to the perfusion protocol and clearing with Cubic^30^, the GPI-anchored FP plasmids tolerate clearing conditions for whole cerebellar imaging with light sheet microscopy with better visualization of axon projections and dendritic arbors than cytoplasmic labeling (Fig. 5). This eliminates the need for antibody staining when coupled with very low Cre plasmid concentrations. Cold temperatures are additionally critical for preserving plasmid supercoiling (Extended Data Fig. 2) as well as endogenous fluorescence during tissue processing. As such, care was taken in the handling temperatures of plasmids, in transcardial perfusions with cold solutions, and in cold handling and storage of extracted brains. To clear in Cubic-L, 37°C is required, so this step was limited as much as possible to preserve endogenous fluorescence. Cubic-R shaking was limited to two days at room temperature to minimize impact on FPs. These workflow modifications enabled us to perform a 3D developmental time-course of generation-tracked Purkinje cells in the mouse brain.

### Insights from stereo-tracking into cerebellar development

Purkinje cells are reported to be orchestrators of cerebellar circuit development as a whole^38,39^, so understanding their migration patterns and developmental transformations informs many cerebellar developmental processes. We therefore followed the progression of generation-tracked Purkinje cells at three time points in the first two weeks of life with 3D cellular imaging of intact cerebella.

*In utero* electroporation allows for specific and sparse labeling of Purkinje cells by targeted electrical pulses to the progenitor zone during their brief window of birth. As demonstrated in Fig. 3b, placement of the positive electrode impacts which cells will be under the strongest part of the electric field, however the posterior cerebellum is consistently more strongly electroporated than the anterior cerebellum. Fig. 5a shows size mismatched cytoplasmic fluorophores across developmental stages to eliminate the confounding factors of Cre expression and observe the layering patterns of Purkinje cells as they migrate postnatally. The large size (10.1 Kb) plasmid is unable to electroporate cells in the anterior cerebellum to a detectable degree, but depending on positive electrode placement, it most strongly electroporates the most dorsal or caudal areas of the posterior cerebellum. The smaller plasmid (5.9 Kb) electroporates both anterior and posterior areas as well as many more neurons than the larger plasmid (5.8% RFP:GFP). The placement along the mediolateral aspect of the progenitor zone determines the extent to which electroporation will occur laterally. Typically, the positive electrode is oriented to the midline, so that the vermis is more heavily electroporated. When the rectangular shaped electrode is placed along the midline, sagittal stripes are electroporated (Fig. 5, P7 movie S2), showing that neurons migrate directly out from their place of birth along the mediolateral and anterior to posterior aspects of the ventricular zone and do not travel to occupy areas away from their birth location. Light sheet imaging allows for examining the topography as a whole.

When Cre recombinase is expressed, the patterns shift depending on plasmid size (8.4 Kb) and concentration (Fig. 5b). The anterior cerebellum is now electroporated because few copies of the Cre plasmid are needed to visualize each cell, but because this area is not as layered as the posterior cerebellum most cells are still at the surface and able to be electroporated with the Cre plasmid, turning them GFP positive. The additional layers of cells deeper into the tissue above the 4^th^ ventricle in the posterior cerebellum are reachable by the smaller fluorophore plasmid (7 Kb) where the larger plasmid does not travel and therefore remain RFP positive. Changing the Cre concentration would determine the ratio of cells electroporated at the surface (Fig. 2). By P7, Purkinje cells have reached their monolayer position so layering based on deep-labeled RFP and superficial-labeled GFP cells are no longer apparent within lobules (Fig. 3a). At P0 (Fig. 5a), the faster GFP plasmid more heavily electroporates deeper into the progenitor zone, reaching cells that have begun to migrate away from the ventricular surface, with the posterior lobules (INF, P0, CE) showing multicellular layering while the anterior lobules (AD, AB) possessing more compact layering (also demonstrated in Fig. 2j at E17.5).

At P0, nascent postmitotic Purkinje cells radiate away from their progenitor zone, much like neurons of the cerebral cortex. As expected, deeper-labeled Purkinje cells are farther along their migratory route toward the Purkinje plate. While late-born, superficial-labeled Pyramidal cells push past early-born, deep-labeled Pyramidal cells in the cerebral cortex to populate different layers of the cortical column, deep-labeled Purkinje cells join superficial-labeled cells in the same monolayer. However, the anterior lobules are overwhelmingly superficial labeled because of the narrower depth-of-field of the anterior progenitor zone at the time of electroporation. The anterior areas of the ventricular zone must also accommodate neurogenesis of cerebellar nucleus interneurons from E10-E12 in the area expressing Gsx1^40^. The anterior and lateral portion of the developing cerebellum appears to have less overall Purkinje cell numbers and multicellular layering within clusters in the anteroposterior plane^26^ and Purkinje cells born on E10.5 are a small subset in the anterior zone, biasing them toward a GFP label^27^. Depending on the Cre concentration and electroporation strength, some layering in the anterior area can be seen (Fig. 2j), though is limited due to the progenitor zone being more compact than in the posterior zone (Fig. 5a P0, Extended Data Fig. 4).

Together, electroporation of the cerebellum reveals that Purkinje cells remain in the general quadrants in which they were born in the ventricular zone, similar to previous reports^4,33^. Cells born along the midline migrate outward to occupy the vermis (P7 movie S2) and cells born more laterally (P14 movie S2) occupy the lateral hemispheres. Anterior-born cells reside in the mature anterior lobules and posterior-born cells reside in the mature posterior lobules. These different quadrants show distinctly different rates of Purkinje cell maturation based on morphology within specific lobules (e.g. Fig. 3c).

The few brightest cells in the vermis are the most heavily electroporated under these conditions, revealing that individual cells are highly stereotyped in their migration patterns. Which subtypes of Purkinje cells (based on heterogeneous expression of genes within clusters) are represented in these specific birth locations remains unclear, but combining the agnostic plasmid expression with stains for specific markers presents an interesting future research direction to track how clusters transform into distinct parasagittal stripes. Since the different stripe markers far outnumber the clusters from which they emerge^7^, it is possible that the deep– and superficial-labeled cells within one cluster, bisected by our stereo-tracking approach, will have differential cascades of gene expression based on the progenitor zone field that commits their differentiation, manifesting in the observed mature stripe patterning. It is thought that compartmentalization begins as early as E10, with the cerebellar stripe arrays developing from the early dorsal to ventral layering in the ventricular zone^8^ so it will be useful to see where electroporated cells overlap with stripe markers using the sparse labeling strategy to more readily track individual cell transformations to their mature locations.

With our improved methods for preserving endogenous fluorescence, we tested whether transgenic lines could similarly be imaged with light sheet for tracking complicated stripe topography, such as that of aldolase C (zebrin II). P90 aldoc-venus mice^41^ highlight the utility of whole cerebellar 3D imaging for examining circuit architecture (Fig. 5c). Purkinje cell patterning based on parasagittal stripe markers is typically represented in 2D from staining many sections. However, observing whole populations in their native circuit architecture allows for examining the continuity of parasagittal stripes across lobules, which was previously unknown for aldolase C, the most widely studied stripe marker^42^. For example, the stripes in the cerebellar hemispheres appears less defined in sections than in the vermis, but in 3D the “ribbons” of aldolase C positive cells connecting across and within lobules can be observed, which raises the question of how signaling across these Purkinje cell types may synchronize to contribute to behaviors. Building accurate models of these topographical features will be important for generating hypotheses on physiological function.

## ADVANTAGES/LIMITATIONS

The technical innovation of stereo-tracking offers a pulse-chase developmental labeling strategy that can be combined with other labeling and analytical strategies for studying the developmental patterns of the brain. It additionally highlights the importance of plasmid handling and size matching when co-electroporation of plasmid mixes is desired to avoid unintended artifacts of labeling. Further technical improvements for labeling include the use of tritrode electroporation for higher efficiency and better targeting of the Purkinje cell progenitor zone; using supercoiled plasmids of small overall size to increase labeling efficiency; and cold-processing brain tissue combined with low Cre plasmid concentrations to preserve endogenous fluorescence for light sheet imaging and to visualize topography and projections in 3D.

Labeling at early embryonic stages can have low survival rates and whether electroporation and expression of plasmids alters the normal development of neurons is unclear. Contrary to what was previously assumed, plasmids do not only electroporate cells directly on the ventricular surface, but also cells that are deeper into the progenitor zone; therefore, cells that are not born on the same day as the electroporation will also be electroporated, which can limit developmental interpretations if this effect is unintended. Particularly in the cerebellum, where patterning of embryonic clusters and mature stripes along multiple planes is informative, electroporation ignores these molecular and anatomical boundaries by pulsing across the entire volume of the ventricular zone and targeting cells at different stages during the 3 days of Purkinje cell birth. Carefully selecting EKM properties of plasmids and electric field orientation can help further identify how distinct fields and depth of cells labeled at the progenitor zone translate into mature patterns of circuitry, both in the cortex and cerebellum.

We document the presence of axon bubbles in stereotyped positions of P14 Purkinje axon initial segments. However, we have not identified neither their function, nor their mysterious internal content. Cyro-EM would help better preserve and interpret membranes of axon bubbles. Serial sections of TEM Purkinje cells would help to determine the precise points of contact along the axon. While we speculate that axon bubbles are a transient feature of developing Purkinje cells due to their tight developmental onset and scattered presence, we have not determined whether they disappear in the mature cerebellum. Future studies will be useful to determine their matrix, function, and defined developmental window, as well as targeted investigations on whether homologous structures may exist in other neuron types in development.

## DISCUSSION

Here we describe a bioelectric interaction of plasmids injected into embryonic progenitor zones and targeted to neural progenitors by pulsed application of an electric field. Delipidated brains have similar properties to an electrolyte gel^43^, and the differential labeling we see is consistent with an electrogenic phenomenon similar to DNA gel electrophoresis. Small, supercoiled plasmids penetrate deeper into the progenitor zone and thus target earlier-born cells on their way to becoming migrating postmitotic neurons. We took advantage of this phenomenon to develop stereo-tracking, a differential labeling strategy that narrowly distinguishes between cells at different depths within the volume of ventricular zone progenitor fields at the developmental pulse point and applied this strategy to investigate how progenitor zone location impacts cerebellar development.

Studies of Purkinje cell migration and rearrangement of clusters heavily rely on known markers at various stages^3,4^. Stereo-tracking allows the tracking of Purkinje cells without regard to subtypes so that the overall topography based on location can be observed and is not reliant on marker staining or transgenic lines. This has been a barrier to studying the development of Purkinje cells, as many markers are not stably expressed throughout development, making it difficult to track cells from the immature cluster stage to mature locations, especially in the significant transitions around P6^4^. Because Purkinje cells born on specific days lead to adult sagittal stripe patterning after the cluster stage^5^, the developmental migration patterns of Purkinje cells appear to be more complicated than other structures. We show with stereo-tracking that Purkinje cells follow a similar overall pattern of migration as in the cerebral cortex at early stages (E17.5-P3) in that migration of cells proceeds directly from the place of origin to the expanding layers above. The cells deeper in the progenitor zone away from the ventricular surface lead the way in pushing the superficial regions of the cerebellar cortex outward, and cells labeled at the superficial fields of the ventricular zone follow closely behind. However, unlike the inside-out layering of the cortex where stereo-tracked cells populate distinct layers, later-born Purkinje cells populate the same monolayer but distinct topographical fields in the mature cerebellar cortex.

Purkinje cell clusters have been reconstructed from 2D images to 3D representations^4^ in an attempt to unravel the seemingly complicated transition to adult stages. However, directly observing structures in 3D without the distortions from tissue processing is an advantage of light sheet microscopy and aids in determining the continuity of structures in the highly foliated cerebellum. Combining 3D imaging with stereo-tracking and other electroporation-based approaches allows for specific targeting of Purkinje cells, as the developmental markers used for identification may also label other cell types and can obscure the topography of pure Purkinje cell populations. Additionally, varying the type of fluorophore tags used in electroporation approaches can aid in highlighting different cellular features, as demonstrated by our unexpected Purkinje cell axon bubble discovery.

## SUPPLEMENTARY DATA

**Extended Data Figures**: Extended Data Figures 1-8.

**Movie S1**: 3D rendering of light sheet imaging of shallow-labeled GPI-GFP (green) and deep-labeled GPI-RFP (magenta) stereo-tracked Purkinje cells in the P14 mouse cerebellum, corresponding to Fig. 5b.

**Movie S2**: 3D rendering of light sheet imaging of shallow-labeled RFP (magenta) and deep-labeled GFP (green) stereo-tracked Purkinje cells in P0, P7, and P14 mouse cerebella, corresponding to Fig. 5a.

**Movie S3**: 3D rendering of light sheet imaging of stripes labeled by Zebrin II-YFP (green) in the adult mouse cerebellum, corresponding to Fig. 5c.

## METHODS

### Animals

Outbred CD1 mice were purchased timed pregnant from Charles River Laboratories. Animals were housed in an animal facility with free access to food and water on a 12/12 h light/dark cycle. All experiments involving animal procedures were approved by the Institutional Care and Use Committees (IACUC) of the University of Maryland School of Medicine and Baylor College of Medicine. Embryonic day 0.5 is the morning after a plug is identified and P0 is the day of pup birth.

### Plasmids

All plasmids were custom designed and constructed with a modified mMoClo system.^44^ Parts were cloned with Bsa1 adapters and assembled into expression constructs with a NEB Golden Gate Assembly Kit (NEB #E1602). Plasmids were transformed into One Shot TOP10 (Thermo Fisher C404006) chemically competent E. Coli and extracted with Zyppy Plasmid Miniprep kits (Zymo Research D4036) to perform diagnostic digests and Sanger sequencing (Azenta Life Sciences). ZymoPURE II Plasmid Midiprep kits (Zymo Research D4200) were used to obtain a high concentration of validated plasmids for use in IUE experiments. Plasmid sizes and open reading frames are provided in the results section, figures and Extended Data fig. 1. All plasmids expressed fluorophores or Cre recombinase from the CAG promoter. Myr-RFP H2B-GFP plasmid used in Fig. 4a was a generous gift by Anna-Katerina Hadjantonakis (MSK).

### *In utero* electroporation

*In utero* electroporation was performed as described in^45^, with the tritrode configuration^9^ to target the Purkinje cell progenitor zone at E11.5. DNA to a maximum total concentration of 4 μg/μl was injected into the 3^rd^ or 4^th^ ventricle until filling the entirety of the 4^th^ ventricle. Six square pulses for 50 ms at 24V were delivered with the positive third electrode placed over the midline of the 4^th^ ventricle. P0 pups were screened for fluorescence under a Leica stereoscope and positive pups were kept with the dam until the appropriate developmental age. As areas electroporated vary between animals, plasmid comparisons are best done within one animal, but a minimum of 2 animals were compared for each separate condition.

### Titrating Cre in EKM plasmid mixtures

With the tritrode electroporation approach, each lateral ventricle can be targeted separately with plasmid mixtures while each hemisphere is targeted with the same electrical field (over the midline of the brain). This eliminates the need to interpret differences across animals and imaging to directly compare changes in fluorophore expression. The mosaic plasmid (2 μg/μL) described in Fig. 1D is co-electroporated with either a low Cre concentration (15 ng/μL) in one ventricle or a high Cre concentration (150 ng/μL) in the other ventricle. Low Cre concentrations preserve endogenous fluorophore expression of both RFP and GFP and can be titrated to produce a 50/50 ratio between colors (Extended Data Fig. 3). However, using excessive Cre concentrations results in a loss of the floxed RFP endogenous expression while the GFP expression is reduced. Although not confirmed, we assume that the loss in GFP expression is a result of recombination of the plasmid to nonfunctional forms from the excessive Cre being produced in most electroporated cells. Higher Cre concentrations will result in a greater likelihood of cells being coelectroporated with both the mosaic plasmid and the Cre plasmid, which leads to more cells that are GFP positive up until the point that plasmids are presumably recombined to non-functional forms. The higher the Cre concentration, the more likely it is to have low overall RFP expression because only the cells subjected to the weakest electrical pulses will receive few copies of the RFP plasmid and not any Cre plasmid.

Weak electroporation results in a lower copy number of plasmids that enter the cells and therefore appear dimmer than cells that receive higher copy numbers under a stronger electrical field.^46^ This can be observed in Extended Data Fig. 3b, where the RFP is strongest in the deeper layers and progressively fades in the upper layers while GFP expression is weak in the deeper layers and bright in the superficial layers. The combination of mosaic plasmid copy numbers that contribute to the fluorophore pool and electroporation of the Cre plasmid based on its size/concentration lead to the overall pattern of expression. Because the ratio of GFP to RFP is sensitive to small changes in Cre concentration, efforts must be made to use the same aliquots of Cre plasmid across experimental animals. Different stocks (as in Extended Data Fig. 3) have to be titrated to achieve the desired ratio of GFP to RFP. When one fluorophore color becomes dim due to low copy numbers of plasmids, expression in distal processes can be lost in comparison to the brighter fluorophore. Additionally, as plasmids lose supercoiling, they effectively have less concentration of supercoiled plasmid that will be electroporated (Extended Data Fig. 2). Therefore, proper storage of plasmids is important and should be run undigested on a gel periodically to detect shifts in supercoiled state.

### Perfusion and tissue processing

Mice were transcardially perfused with ice cold PBS followed by 4% PFA and postfixed in cold 4% PFA for 24 hours. Sections were sliced on a vibratome in cold cutting conditions at 80 μm, mounted on glass slides and coverslipped with Fluoromount mounting media (Thermo Fisher 00-4958-02) then imaged at 10x (Nikon Ti2-E inverted microscope) immediately for best retention of endogenous plasmid expression. Room temperature sections (in PBS) lose fluorescence intensity over a few hours, so must be kept cold, otherwise will need to be stained with immunofluorescence. Whole cerebellum images were taken with a Leica stereoscope directly after perfusion and before further processing to avoid high GFP background after post-fixation.

### Clearing and light sheet imaging

Brains were cleared to transparency as described^47^ by a modified Cubic protocol.^30^ Whole PFA-fixed brains were placed into 100% Cubic-L in a shaking water bath at 37 °C. Cubic-L was replaced every two days until the brain was uniformly opaque and white matter tracts were no longer seen (about one week). Brains are then briefly rinsed in Cubic-R+(M) and placed in fresh Cubic-R+(M) shaking at room temperature until transparent (about two days). Whole brain images were acquired with a Zeiss Light Sheet 7 with lenses adjusted to a refractive index of 1.52 with a 5x objective and visualized with Arivis software. Arivis segmentation of individual Purkinje cells was achieved with the “blob finder” analysis pipeline. Each identified cell is listed as an object, all objects found with each color (RFP and GFP) were counted then RFP was expressed as a percentage of GFP expressing cells.

### Transmission electron microscopy

Purkinje cells from the cerebellum were subjected to TEM imaging following established protocols. After transcardial perfusion of Ringer’s solution followed by modified Karnovski’s fixative (pH 7.4), sagittal sections of the cerebellum measuring 1-2 mm in thickness were prepared at room temperature. These sections were then immersed in modified Karnovski’s fixative in 0.1 M sodium cacodylate buffer at pH 7.2 and rotated in scintillation vials for a period of three days. On the third day, the tissue was processed inside a Ted Pella Bio Wave Vacuum Microwave on ice. Samples were fixed again, followed by 3x sodium cacodylate buffer rinses, post-fixed with 1% buffered osmium tetroxide, and followed again with 3 millipore water rinses. Ethanol concentrations from 30-100% were used as the initial dehydration series, followed with propylene oxide as a final dehydrant. The samples were gradually infiltrated with propylene oxide and Embed 812 resin, with several changes of pure resin under vacuum. Following overnight infiltration in pure resin on a rotator, the samples were embedded into regular Beem capsules and cured in an oven at 62 °C for five days. After polymerization, the embedded samples were thin-sectioned to 50 nm and stained with 1% uranyl acetate for fifteen minutes, followed by lead citrate for two minutes before TEM examination. TEM imaging was performed using a JEOL JEM 1010 transmission electron microscope with an AMT XR-16 mid-mount 16 mega-pixel digital camera. Subsequent to imaging, adjustments to image contrast were made using ImageJ software.

## ACKNOWLEDGMENTS

We are grateful to Bekir Altas, Colin Robertson (Baltimore), and Andy Cole (NIH) for very useful discussions, and to Joseph Mauban and Thomas Blanpied for the technical and instrumentation support at the University of Maryland School of Medicine Center for Innovative Biomedical Resources through the Confocal Imaging Facility and grant S10 OD030221 (Zeiss Light sheet) and Lita Duraine for her expertise in preparing samples and imaging with TEM. CB is exceptionally grateful to Dr. Greg Elmer for his support and mentorship throughout the project under the T32 fellowship. We thank Apollo, Sun God, for the spring equinox when light overcomes dark and dreams awake into realities.

## AUTHOR CONTRIBUTIONS

Conceptualization: CB, GWC, AP; Methodology: CB; Validation: GWC, AJR; Formal Analysis: CB, AP Investigation: CB, SGD, LRD; Resources: O.-M.R.N.A, IS, RVS, AP; Writing-Original Draft: CB; Writing-Review and Editing: CB, GWC, AJR, SGD, LRD, O.-M.R.N.A, IS, GJB, RVS, AP; Visualization: CB, LRD, AP; Supervision: RVS, AP

