## Extended Data Figure for "Developmental transformations of Purkinje cells tracked by DNA electrokinetic mobility"

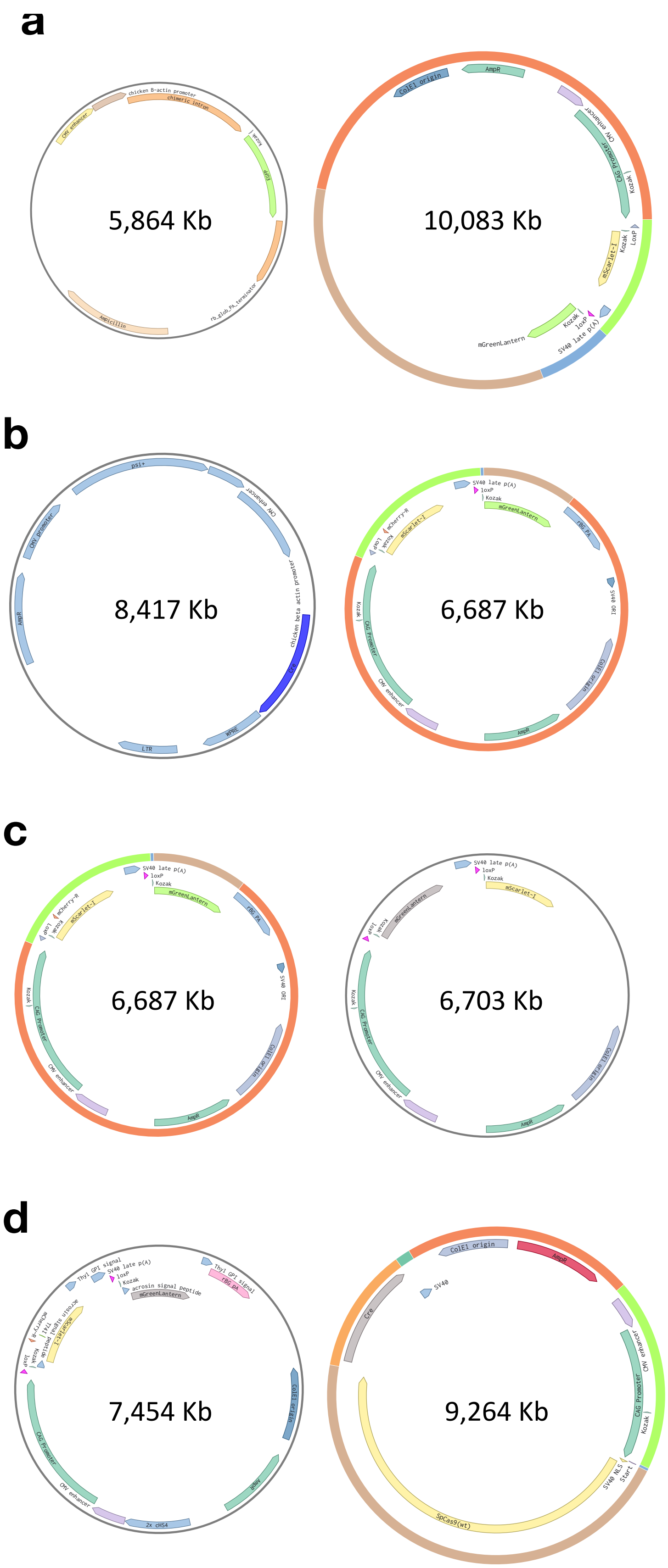


**Extended Data Figure 1. Construct maps and sizes of plasmid pairs used to determine the effect of electrokinetic mobility on *in utero* electroporation.**

**a,** Fig. 1b,c; Fig. 5a **b,** Fig. 1e; Fig. 2, Fig. 3b **c,** Fig. 1d **d,** Fig. 5b. All pairs aside from **c** are unmatched in electrokinetic mobility. Inserts located after poly A tails will not be translated so can be used to increase plasmid size without transcriptional impact. Backbones elements (as in **b**) were removed without expression consequences to decrease size and match the partner plasmid's size.

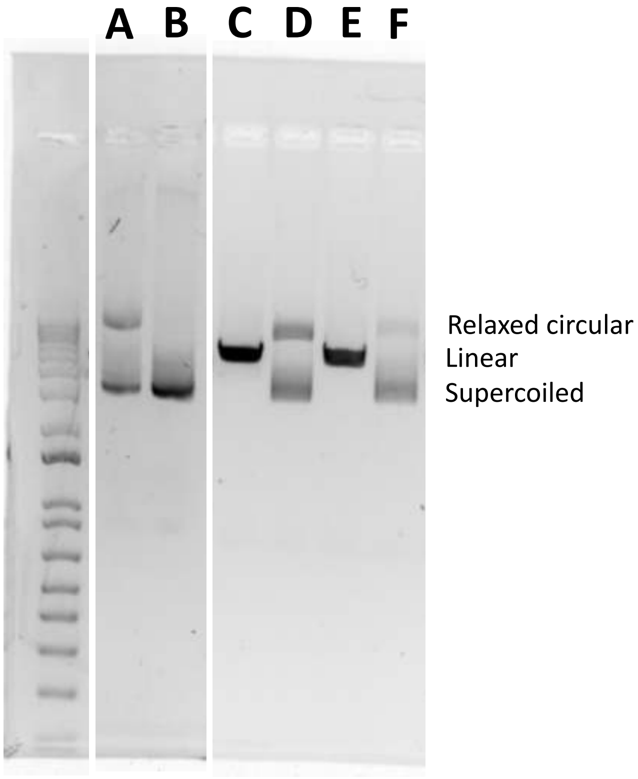


**Extended Data Figure 2. Plasmid elektrokinetic mobility changes in gel electrophoresis.** Each lane (A-F) represents the same plasmid of size 5,956 Kb, run on two different gels. Lane A was a several month old plasmid prep subjected to periods of room temperature and freeze-thaw cycles, while lane B was the same plasmid freshly prepared and frozen at -20℃ (run undigested). Lane B does not show the open circular form present in Lane A from extended storage times at higher temperatures (Shleef et al., 2006) and is entirely in the supercoiled form. The plasmid prep in lane A is poorly transfected during in utero electroporation while the lane B plasmid has high transfection rates. The different preps from lane A and B both show the same linear band when a restriction digest is performed to cut the plasmid in one location (lanes C and E). When incubated at 37℃ for one hour and then run undigested, the compromised plasmid (from lane A) further deteriorates away from the supercoiled form and toward the relaxed circular form (lane D), while the fresh prep from lane B also begins to show relaxation, but to a lesser degree (lane F). Plasmids used for in utero electroporation should be run undigested to ensure all preps maintain most of their concentration in the supercoiled form.

Schleef, M., Baier, R., Walther, W., Michel, M.-L., & Schmeer, M. (2006). Long-Term Stability Study and Topology Analysis of Plasmid DNA By Capillary Gel Electrophoresis. BioProcess Int, 4.

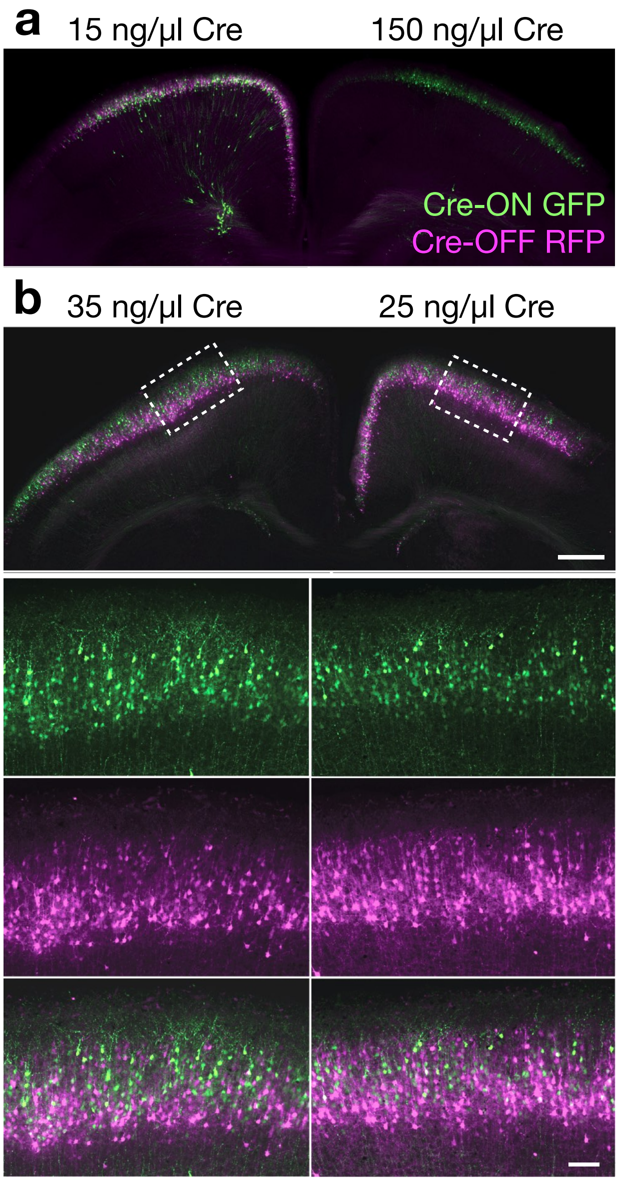


**Extended Data Figure 3. Titrated EKM plasmid mixtures enable "progenitor zone stereo-tracking" by differentially targeting distinct depths of the the progenitor field with one electroporation.**

Each lateral ventricle in the same animal was separately injected with the Cre-dependent mosaic plasmids (plasmids b in Extended Data Fig. 1) expressing Cre-ON GFP (green) at two different concentrations of Cre plasmid (ng/µl plasmid DNA as indicated), differences in expression of the fluorophores in neuron populations and fluorescent protein intensity. **a,** Electroporation of embryos at E16.5 suppresses layer segregation since no further cortical neuron layers are produced during beyond that time point. The hemisphere that received 150 ng/µl Cre plasmid showed diminished RFP expression, indicating that most cells received Cre, which excised the RFP open reading frame. **b,** Layered labeling appear electroporation at E14.5 and earlier. Cre concentration effects layering segregation as well as and the ratio of GFP to RFP expressing cells. Cells at the ventricular surface during electroporation and migrating to superficial layers receive higher concentrations of Cre (GFP positive), while the slow Cre plasmid does not reach as many cells away from the ventricular surface at the time of electroporation, which migrate to deeper cortical layers (RFP positive). Scale bar 500 µm in main panel, 100 µm in inset.

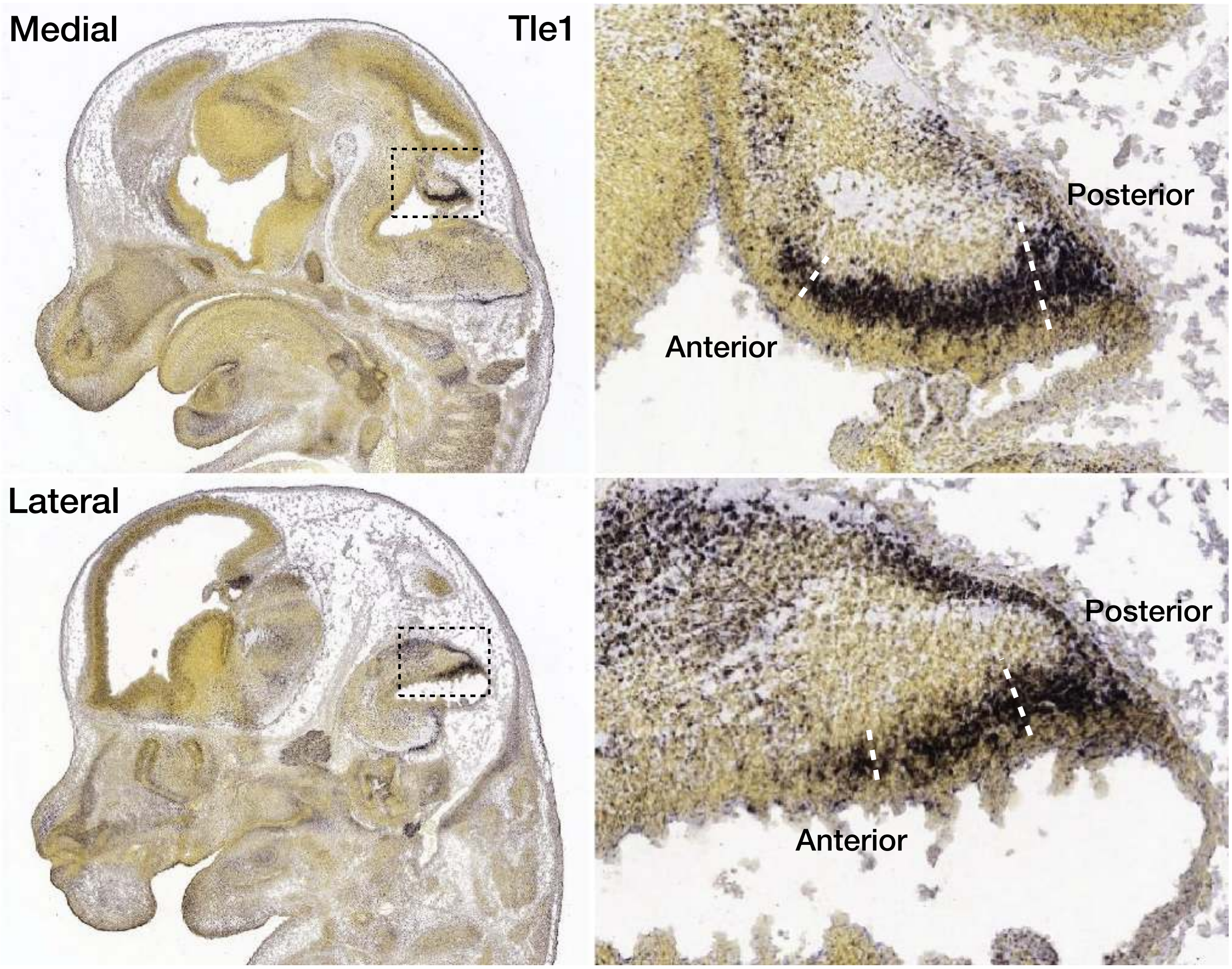


**Extended Data Figure 4. The Purkinje cell progenitor zone.** Allen Developing Brain Atlas labeling all Purkinje cells at E13.5 as they form the Purkinje cell plate. In the cerebellum, *Tle1* is a Purkinje cell marker in early development (Wizeman et al., 2019) and serial sections are provided by the Allen Institute after *in situ* hybridization with *Tle1* probe. In the medial posterior, Purkinje cell layering is thickest, which eventually becomes the vermis that is labeled most densely with RFP (Fig. 2 in main). Moving both in the anterior and lateral direction, the Purkinje cell field becomes thinner, with the least number of Purkinje cell layers across the anterior portion of the ventricular zone (in the dorsal to ventral plane of layering). A larger proportion anterior cells are close to the ventricular surface at the time of the electroporation thus mainly become GFP expressing (Fig. 2 in main).

Wiseman, J.W., Guo, Q., Wilson, E.M. & Li, J.Y.H. Specification of diverse cell types during early neurogenesis of the mouse cerebellum. Life **8**, 1-24 (2019).

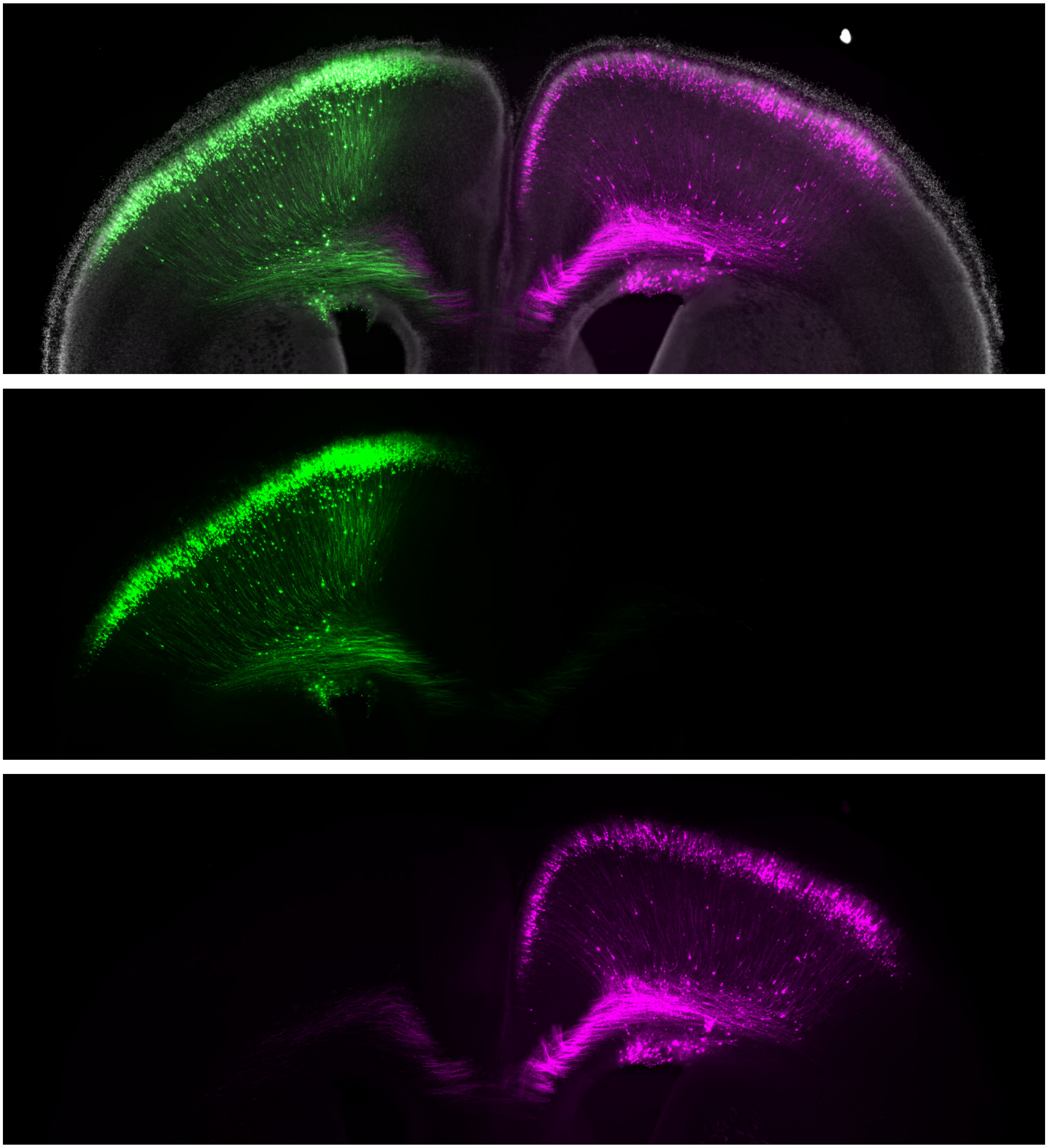

**Extended Data Figure 5. Dual Electroporation.** Bilateral plasmid injections into the ventricles stay segregated to each cortical hemisphere during in utero electroporation (shown at P3). The plasmid pair in Extended Data Fig. 1c was injected into separate ventricles without a Cre plasmid present, showing that the polyA tail is effective at preventing translation of the second fluorophore.

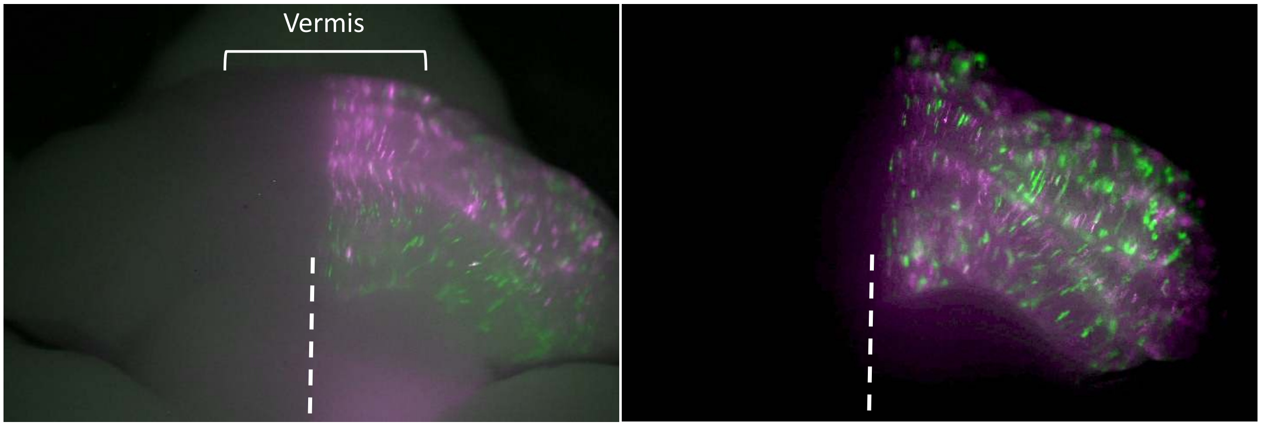


**Extended Data Figure 6. Stereo-tracking versus balanced EKM electroporations in the cerebellum.** Size-matched plasmids equally distribute in the cerebellum. Compared to Figure 3C (left image) of the main text, the bright area of RFP expression (magenta) in the vermis is eliminated (right image) and the anterior lobules now have equal representation of RFP and GFP (green).

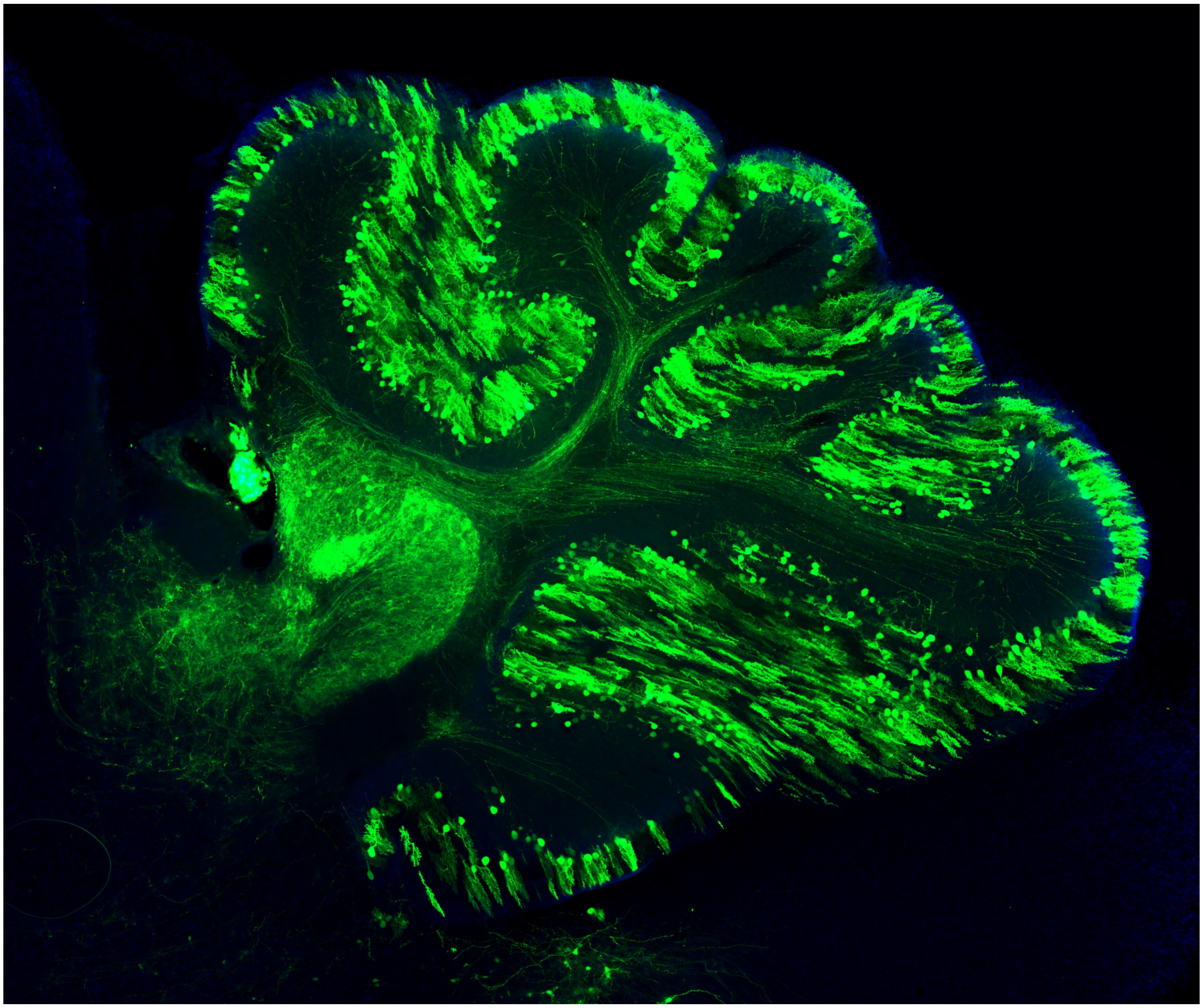

**Extended Data Figure 7. Cerebellar section at P14 showing Purkinje cells expressing cytosolic GFP.** Cytosolic fluorophores more brightly label cell bodies. Simultaneously imaging both axons and cell bodies results in saturated cell bodies but dim axons. This presents a particular obstacle with 3D light sheet imaging where bright cell body fluorescence scatters in the unlabeled cleared tissue increasing background. Additionally, cytosolic FPs do not label axon bubbles.

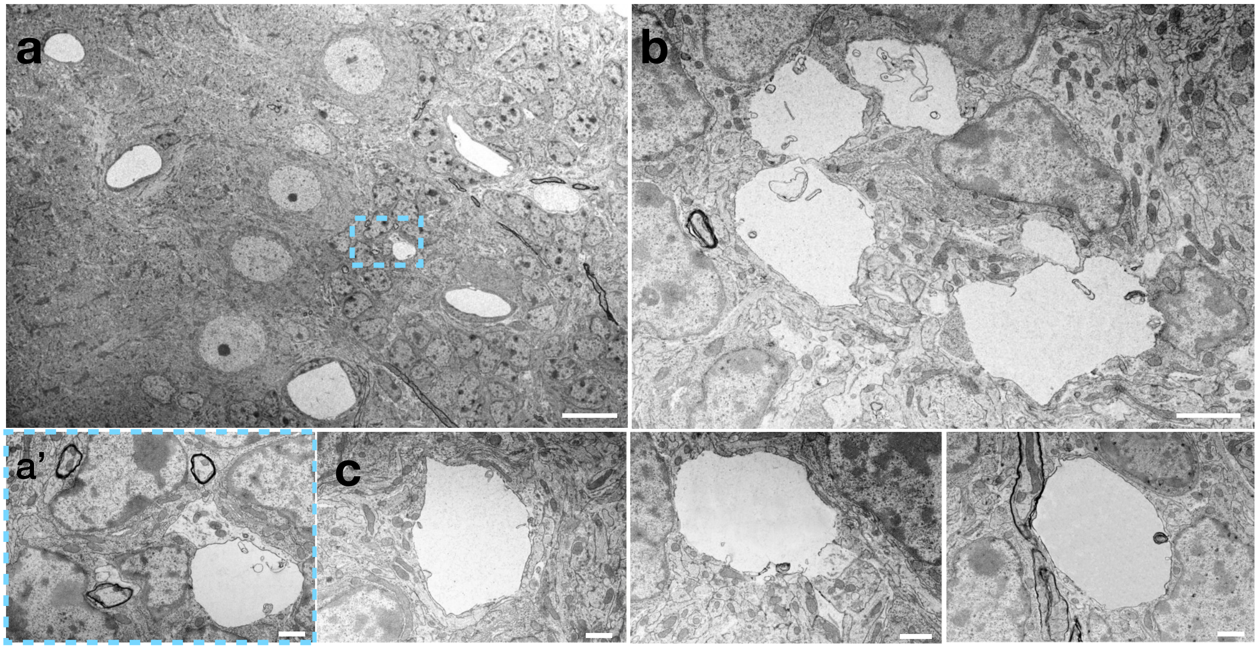


**Extended Data Figure 8. Electrolucent bubbles on Purkinje cell axon initial segments. a,** Low electron micrographs of Purkinje cell layer showing, in addition to Figure 5 of the main text, that electrolucent axon bubbles appear sporadically about 10 µm from the Purkinje cell soma on the granule cell layer side, which are unlike blood vessels seen throughout the tissue. The area outlined in dashed blue, containing the apple-shaped axon bubble, is magnified in α’. Scale bars 10 µm, 1 µm inset. **b-c**) Higher magnification of individual axon bubbles highlights their irregular shapes, lack of endothelial cells/pericytes that differentiate them from blood vessels, and position along structures associated with the axon. Scale bars 10 µm (b) and 2 µm (c).
